# Contributions of h- and Na^+^/K^+^ pump currents to the generation of episodic and continuous rhythmic activities

**DOI:** 10.1101/2020.09.08.288266

**Authors:** Simon A. Sharples, Jessica Parker, Alex Vargas, Adam P. Lognon, Ning Cheng, Leanne Young, Anchita Shonak, Gennady S. Cymbalyuk, Patrick J. Whelan

**Affiliations:** School of Psychology and Neuroscience, University of St Andrews, Fife, United Kingdom; Hotchkiss Brain Institute, University of Calgary, Calgary, AB, Canada; Department of Neuroscience, University of Calgary, Calgary, AB, Canada; Neuroscience Institute, Georgia State University, Atlanta, GA, USA; Department of Comparative Biology and Experimental Medicine, University of Calgary, Calgary, AB, Canada; Department of Physics and Astronomy, Georgia State University, Atlanta, GA, USA

**Keywords:** episodic rhythms, central pattern generator, spinal cord, rhythmicity, dopamine

## Abstract

Developing spinal motor networks produce a diverse array of outputs, including episodic and continuous patterns of rhythmic activity. Variation in excitability state and neuromodulatory tone can facilitate transitions between episodic and continuous rhythms; however, the intrinsic mechanisms that govern these rhythms and their transitions are poorly understood. Here, we tested the capacity of a single central pattern generator (CPG) circuit with tunable properties to generate multiple outputs. To address this, we deployed a computational model composed of an inhibitory half-centre oscillator (HCO). Following predictions of our computational model, we tested the contributions of key properties to the generation of an episodic rhythm produced by isolated spinal cords of the newborn mouse. The model recapitulates the diverse state-dependent rhythms evoked by dopamine. In the model, episodic bursting depended predominantly on the endogenous oscillatory properties of neurons, with Na^+^/K^+^ ATPase pump (I_Pump_) and hyperpolarization-activated currents (I_h_) playing key roles. Modulation of either I_PumpMax_ or I_h_ produced transitions between episodic and continuous rhythms and silence. As I_Pump_ increased, the episode duration and period increased along with a reduction in interepisode interval. Increasing I_h_ increased the episode period along with an increase in episode duration. Pharmacological manipulations of I_h_ with ZD7288 and I_Pump_ with ouabain or monensin in isolated spinal cords produced findings consistent with the model. Our modelling and experimental results highlight key roles of I_h_ and I_Pump_ in producing episodic rhythms and provide insight into mechanisms that permit a single CPG to produce multiple patterns of rhythmicity.

**Significance statement:** The ability of a single CPG to produce and transition between multiple rhythmic patterns of activity is poorly understood. We deployed a complementary computational half-centre oscillator model and an isolated spinal cord experimental preparation to identify key currents whose interaction produced episodic and continuous rhythmic activity. Together, our experimental and modelling approaches suggest mechanisms in spinal networks that govern diverse rhythms and transitions between them. This work sheds light on the ability of a single CPG to produce episodic bouts observed in behavioural and pathological contexts.

## Introduction

Locomotor behaviours enable organisms to navigate and interact with their environment. These behaviours are remarkably diverse and flexible, but many species share a defined set of locomotor acts that are often engaged in specific contexts. For example, locomotor behaviour during foraging is often episodic, interspersed with pauses to survey the environment. In contrast, pauses are less favourable during migratory locomotor behaviours, which are generally continuous to maximize distance travelled. Animals also need the capacity to rapidly switch between locomotor behaviours. When foraging animals sense a predatory threat, they need to respond quickly by initiating a freeze or escape response (Kim et al. 2017; Ferreira-Pinto et al. 2018). In mammals, the neural mechanisms that govern the generation of episodic and continuous locomotor bursts and transitions between them are unclear, but studies in fish and *Xenopus* tadpoles provide evidence that an interaction between descending commands, sensory modulation, and endocannabinoids contribute (Berg et al. 2018; Eaton, Lee, and Foreman 2001; Sillar and Robertson 2009).

Mounting evidence suggests that the spinal central pattern generator (CPG) encodes episodic and continuous locomotor patterns. Episodic and continuous patterns of rhythmicity can be generated in vitro in neonatal mouse, zebrafish, and *Xenopus* tadpole spinal cord preparations by varying their neuromodulatory tone and excitability state (Picton, Sillar, and Zhang 2018; Sharples and Whelan 2017; Mahrous and Elbasiouny 2018; Kondratskaya et al. 2019; Montgomery et al. 2021; Wahlstrom-Helgren et al. 2019; Wiggin et al. 2012; McDearmid and Drapeau 2006; Marchetti and Nistri 2001; Gozal et al. 2014). In larval zebrafish, features of episodic locomotor behaviours change during development from long, sporadic swimming episodes to shorter episodes with a more regular beat-and-glide pattern (Lambert, Bonkowsky, and Masino 2012; Buss and Drapeau 2001). This switch, which coincides with the onset of foraging behaviours (Borla et al. 2002), is mediated by D_z_dopamine receptors (Lambert, Bonkowsky, and Masino 2012), and patterns continue to mature through juvenile and adult stages of development (Gabriel et al. 2011; Müller, Stamhuis, and Videler 2000). A similar developmental switch in locomotor behaviour has been reported in *Xenopus* tadpoles (Currie et al. 2016). During early developmental stages, D_2_ dopamine receptors promote a sessile behaviour, and at later stages, D_1_ receptors facilitate episodic swimming for filter-feeding (Clemens et al. 2012; Picton and Sillar 2016). Swimming episodes in tadpoles have been linked to the activity-dependent Na^+^/K^+^ ATPase pump current (I_Pump_; (Pulver and Griffith, 2010; Zhang and Sillar 2012; Zhang et al. 2015), which interacts with hyperpolarization-activated cation currents (I_h_) and A-type K^+^ currents (I_KA_) in excitatory rhythm-generating interneurons (Picton, Sillar, and Zhang 2018; Zhang et al. 2015).

The circuit dynamics that produce episodic and continuous network outputs and the mechanisms that drive transitions between these patterns are less clear. In many species, diverse outputs from motor circuits are produced through a combination of dedicated and multifunctional circuit elements that possess tunable properties (Getting 1989; Berkinblit et al. 1978; Mackie and Meech 1985; Meech and Thomas 1977; Gutierrez and Marder 2014; Gutierrez, O’Leary, and Marder 2013; Parker et al. 2018; Li et al. 2007; Ramirez and Pearson 1988; Berkowitz 2002). In support of a multifunctional circuit concept, we previously described a dynamic mechanism and developed a model of a single half-centre oscillator (HCO) that could produce two rhythmic patterns with very distinct cycle periods (CPs): a slow locomotor-like pattern (∼ 1 s) and a fast paw-shake-like pattern (∼ 0.1 s) (Parker et al. 2018; Bondy et al. 2016).

Here, we explored cellular dynamics that enable a single CPG to produce very slow episodic patterns of rhythmicity (episode period [EP] ∼ 50 seconds) and identified changes in intrinsic properties that elicit transitions to faster continuous rhythmicity (CP ∼ 1 s). We adapted our established biophysical model (Parker et al. 2018) to produce episodic and continuous patterns of rhythmic activity, similar to patterns previously observed in the isolated neonatal rodent spinal cord in vitro (Barbieri and Nistri 2001; Sharples and Whelan 2017; Marchetti and Nistri 2001; Gozal et al. 2014). We then compared the contribution of key intrinsic properties predicted by the model to the generation of episodic rhythmicity elicited by dopamine in experiments on isolated neonatal mouse spinal cords. This preparation offers a convenient means to study these dynamics because both episodic and continuous patterns can be elicited (Sharples and Whelan 2017), and its cellular properties can be readily manipulated. Our complementary approaches provide novel insights into how CPGs produce multiple rhythmic patterns and transitions between them. Some of these results have been presented previously in abstract form (Vargas A & Cymbalyuk G 2019).

## Results

### A biophysical model that generates episodic rhythmicity

We developed a biophysical model consisting of two neurons representing mutually inhibiting neuronal populations, an HCO (Figure 1). Our goal was to elucidate a mechanism that explains how a single HCO could generate episodic and continuous bursting, and then compare responses of the model to variations of specific properties with experimental data. We suggest that the dynamics of episodic bursting in the neonatal mouse locomotor CPG rely on the interaction of persistent Na^+^ current (I_NaP_), I_Pump_, and two different types of hyperpolarization-activated current (I_h-a_ and I_h-b_). We started with a model which included only one h-current, I_h-a_, and was capable of producing episodic bursting activity, however, we found that an additional h-current, I_h-b_, was needed to better replicate experimental results of blocking I_h_ with ZD7288. I_h-a_ and I_h-b_ in our model differ mainly by their voltages of half-activation, where the V_1/2 h-b_ (similar to HCN1 and HCN3) is more hyperpolarized than V_1/2 h-a_ (similar to HCN2 and HCN4) (Emery, Young, and McNaughton 2012).

**Figure 1:**
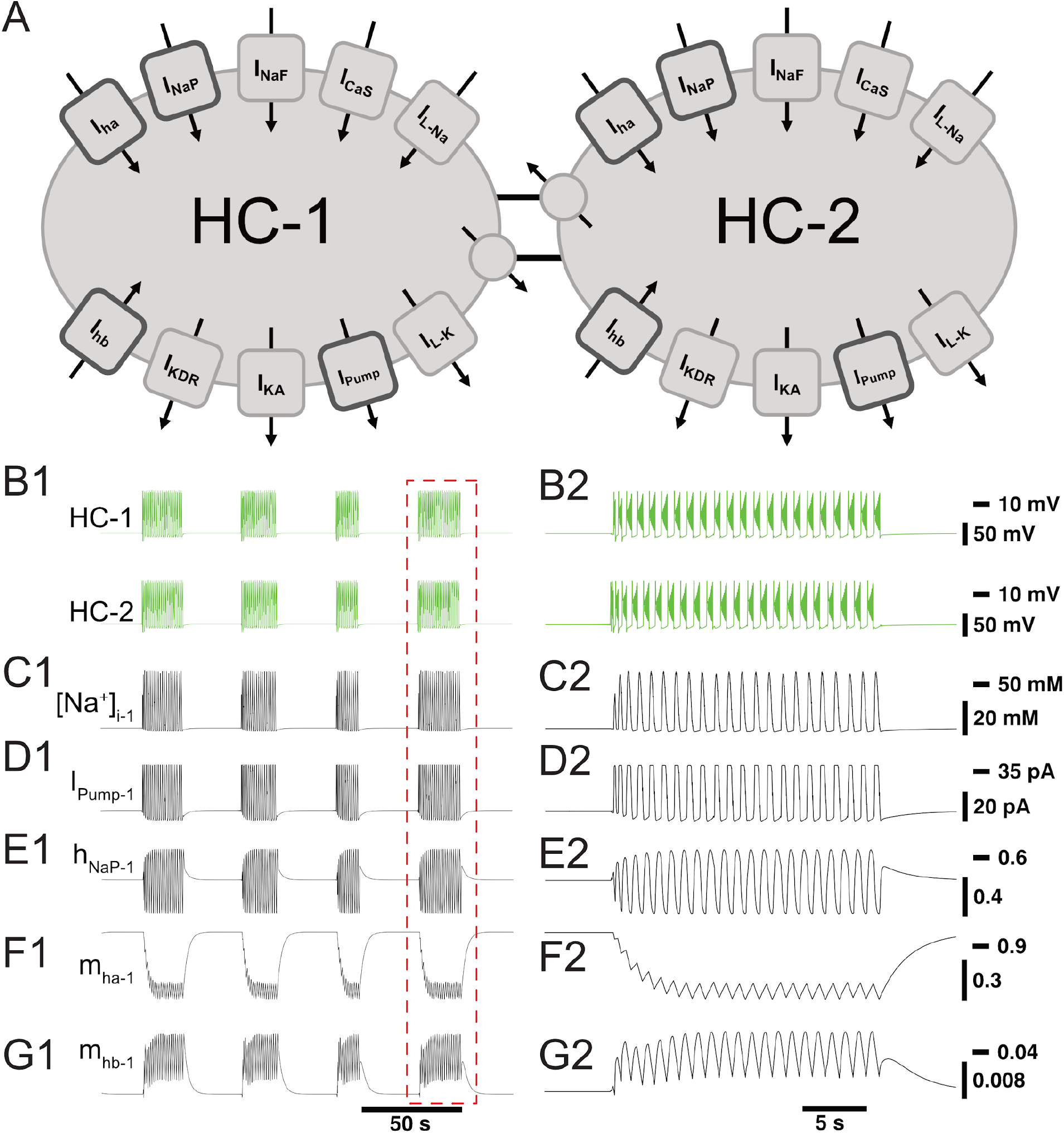
Activation and deactivation of I_h_ governs episodic bursting in a computational model. (A) Schematic representation of the model showing all membrane currents for the two mutually inhibitory cells of the half-centre oscillator (HCO). Inward currents include the slow calcium current (I_CaS_), fast Na^+^ current (I_NaF_), persistent Na^+^ current (I_NaP_), Na^+^ leak current (I_L-Na_), and two hyperpolarization-activated cation current (I_h-a_ and I_h-b_). Outward currents are the potassium leak current (I_L-K_), delayed rectifier potassium current (I_KDR_), A-type potassium current (I_KA_), Na^+^/K^+^ pump current (I_Pump_), and inhibitory synaptic current (I_Syn_). (B) The membrane potential of the two model cells (HC-1 and HC-2) engage in episodic bursting activity (B1) and burst in alternation within an episode (B2). The model is entirely symmetric, so the time course of a few important state variables and currents are shown for only one neuron (neuron 1): (C1-C2) intracellular Na^+^ concentration ([Na]^+^_i1_); (D1-D2) I_Pump1_; (E1-E2) inactivation of persistent Na^+^ current (h_NaP1_); (F1-F2) activation of I_h-a_ (m_h-a1_); and (G1-G2) activation of I_h-b_ (m_h-b1_). The mechanism producing episodic bursting relies on I_h-a_, which slowly deactivates during the episode and promotes its termination (F2).

We validated predictions from the model using an isolated spinal cord preparation, in which we elicited episodic rhythmicity by bath application of dopamine (Figure 2) (Sharples and Whelan 2017). This preparation also allowed us to manipulate and assess the relative contribution of certain currents, which the model predicts play a key role in the rhythmic network activity. Similar forms of episodic rhythmicity are not exclusive to dopamine application (Sharples and Whelan 2017; Sharples et al. 2020); they can be evoked by trace amines (Gozal et al. 2014) and activation of excitatory tachykinin receptors (Barbieri and Nistri 2001; Marchetti and Nistri 2001) in neonatal mouse and rat spinal cord. Further, episodic rhythms generated in vitro display consistent temporal properties to swim episodes in mature zebrafish (Müller, Stamhuis, and Videler 2000; Gabriel et al. 2011; Currie et al. 2016) and juvenile (P11-15) mouse isolated spinal preparations (Mahrous and Elbasiouny 2018). The mechanism utilized by our model to produce episodic bursting could potentially apply to the generation of episodic bursting in these other organisms as well.

**Figure 2:**
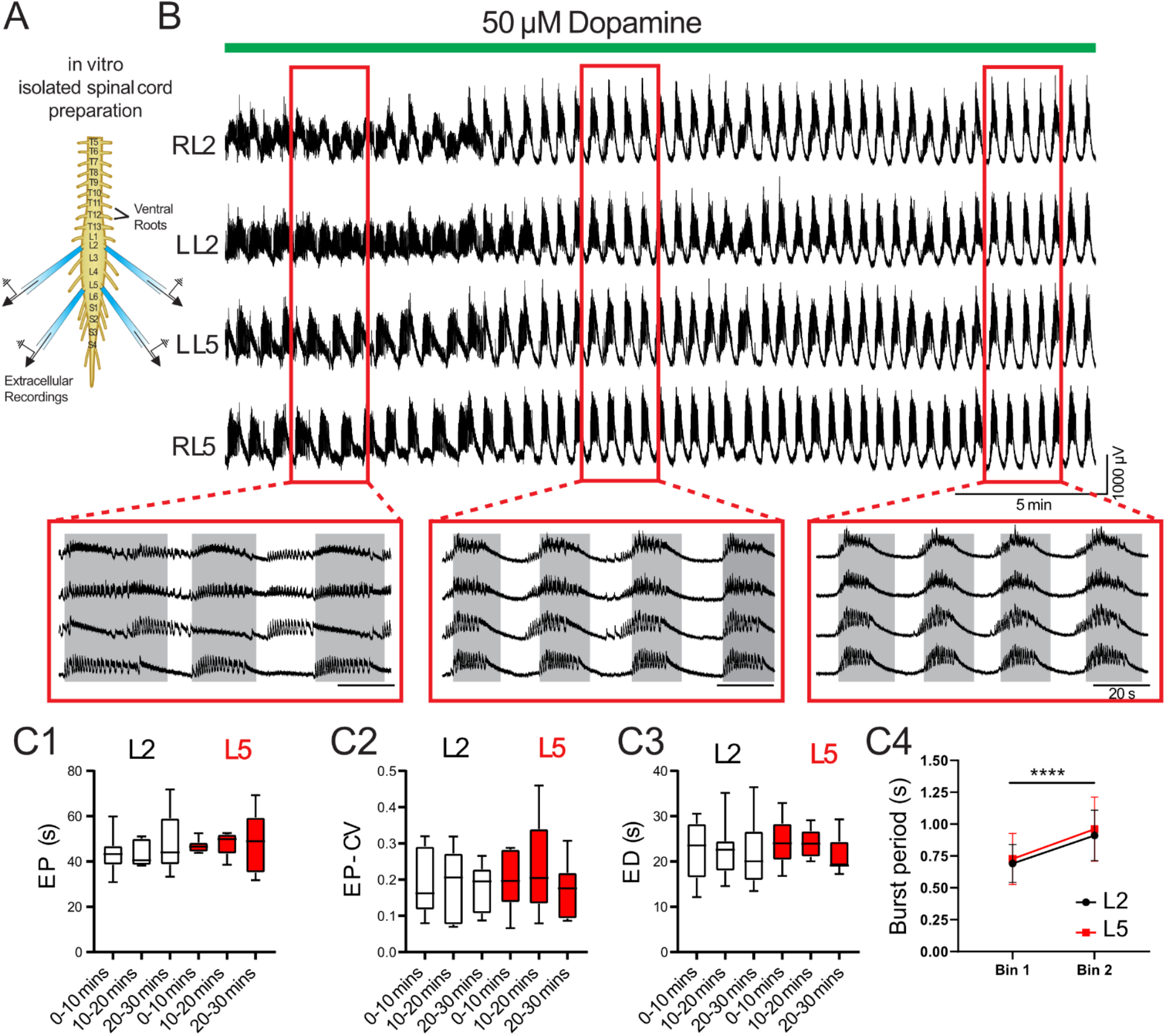
Detailed analysis of dopamine-evoked rhythmicity. (A) Suction electrodes were used to record extracellular neurograms from left (L) and right (R) ventral roots of the second (L2) and fifth (L5) lumbar segments. (B) Dopamine applied at 50 µM (green bar) evoked episodic rhythmicity in lumbar spinal circuits. Expanded red boxes highlight changes in episode patterning across ventral roots over time. (C) From 0-30 minutes after dopamine application, episodes did not differ between L2 and L5 in episode period (EP: C1), episode period coefficient of variation (EP-CV: C2) or episode duration (ED: C3). Box and whisker plots display interquartile range (boxes), median (horizontal black lines), maximum, and minimum values in data range (whiskers). (C4) The burst period of the intra-episode rhythm increased between the first (bin 1) and second (bin 2) halves of an episode. Data in C4 are presented as mean ± SD during the first (Bin 1) and second (Bin 2) half of each episode. Asterisks denote significance (*** p < 0.001, **** p<0.0001) from post hoc analyses following a 2-way ANOVA.

As previously reported, 50 µM dopamine elicited stable intervals of episodic activity with episode period (EP) = 48.3 ± 8.7 s, EP variability defined by the coefficient of variation (EP-CV) = 0.17 ± 0.19, episode duration (ED) = 21.5 ± 19.9 s, and interepisode interval (IEI) = 17.6 ± 15.3 s. Episodes in the L2 and L5 did not differ or vary over time in terms of EP (Figure 2C1; F_(2,14)_=3.4, p=0.11) EP-CV (Figure 2C2; F_(2,14)_= 0.7, p = 0.7), or ED (Figure 2C3; F_(2,14)_=0.28, p=0.6). Consistent with previous reports (Sharples and Whelan 2017), intra-episode bursting slowed over the duration of an episode and did not differ between L2 and L5 (Figure 2D4; F_(1,7)_=0.58, p=0.5).

### Key currents for producing episodic rhythmicity

In our model, the episodic bursting was governed by the slow dynamics of the activation variable of I_h-a_ (m_h-a_) (Figure 1G1, G2). During the IEI, I_h-a_ activated relatively slowly in accordance with its voltage dependence and balance of inward and outward currents. I_h-a_ carried a small amount of Na^+^ into the model cells (Figure 1G1, G2); and this increase in intracellular Na^+^ concentration caused I_Pump_ to increase slightly throughout the IEI. The simultaneous increase in I_h-a_ and I_Pump_ led to subthreshold oscillations in the membrane potentials of both cells. These membrane potential oscillations increased in amplitude over time until the oscillation peak was great enough to activate I_NaP_ and low-threshold slowly-inactivating Ca^++^ current (I_CaS_); they reached an amplitude of approximately 2 mV before the beginning of the bursting phase, characterized by alternating bursting between HCO neurons (Figure 1). Throughout the intra-episode, bursting phase, I_h-a_ slowly deactivated, deactivating further with each consecutive burst for the first 7-10 bursts until it plateaued and oscillated around m_h-a_ = 0.57 (Figure F1, F2). Once this plateau was reached, bursting activity appeared periodic and stable. At this stage, the oscillation properties (such as maximum and minimum) of all of the important variables exhibited slight variability from burst to burst (Fig 1B2-G2). Interestingly, any of these variations could cause the episode to end and fall into the silent phase. Therefore, there was a high degree of variability in the model’s ED.

We empirically identified ranges of parameters around their canonical values (one at a time) within which our HCO model exhibited episodic activity. Changing I_Pump_ and I_h-b_ moved the system into and out of a zone supporting episodic bursting. The hyperpolarization-activated currents, I_h-b_ and I_h-a_, are mixed ion currents carried by Na^+^ and K^+^ ions and the activation of I_h-a_ (m_h-a_) was the slowest state variable of the system (Figure 1G1, G2). Thus, in addition to their roles as predominantly inward currents, I_h-a_ and I_h-b_ also contributed to intracellular Na^+^ concentration ([Na^+^]_i_), which was the second slowest variable of the system. In each case of the specific parameter variation, we compared changes predicted by the model with experimental data from the isolated spinal cord preparation. We found that the parameters of I_Pump_, I_h-a_, I_h-b_, and the inhibitory synaptic current (I_Syn_) required careful adjustment to fit the temporal characteristics of experimental data. The burst generating mechanism of our model was based on the dynamics of activation and inactivation of I_NaP_ and I_CaS_, which controlled the initiation and termination of individual bursts within an episode. I_NaP_ and I_CaS_ also contributed to episode termination along with I_h-a_, I_h-b_, and I_Pump_. I_h-a_ contributed to episode characteristics by controlling ED and IEI, while I_h-b_ primarily affected ED.

### Model-generated episodic rhythmicity is consistent with spinal rhythms generated in vitro

We tuned maximal pump current (I_PumpMax_) and the maximal conductance for I_h-b_ 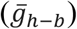 of the model such that it exhibited episodic bursting activity with temporal characteristics close to episodes generated by isolated spinal cords in response to 50 µM dopamine (Figure 2; Figure 3A, B). Using the parameters I_PumpMax_ = 39.9 pA and 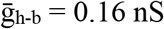 in our “canonical” parameter set, episodes generated by the model were not significantly different from those recorded in vitro from the spinal cord (n = 8) in terms of EP (Figure 3D1, *t*_(63)_= 0.05, *p* = 0.96 s), EP variability (EP-CV; Figure 3D2, *t*_(7)_ = 0.1, *p* = 0.92), or ED (*t*_(64)_ = 0.9, *p* = 0.4; Figure 3D3). Furthermore, the model could reproduce intra-episode burst period dynamics observed in vitro, with an intra-episode burst period that started short and increased in length during the episode (Figure 3C1, C2). Intra-episode bursting generated by the model, which started with a burst period of 0.36 s and increased to 1.0 s by the end of the episode, differed from experimental data that started with a burst period of 0.65 ± 0.2 s and increased to 0.9 ± 0.2 s.

**Figure 3:**
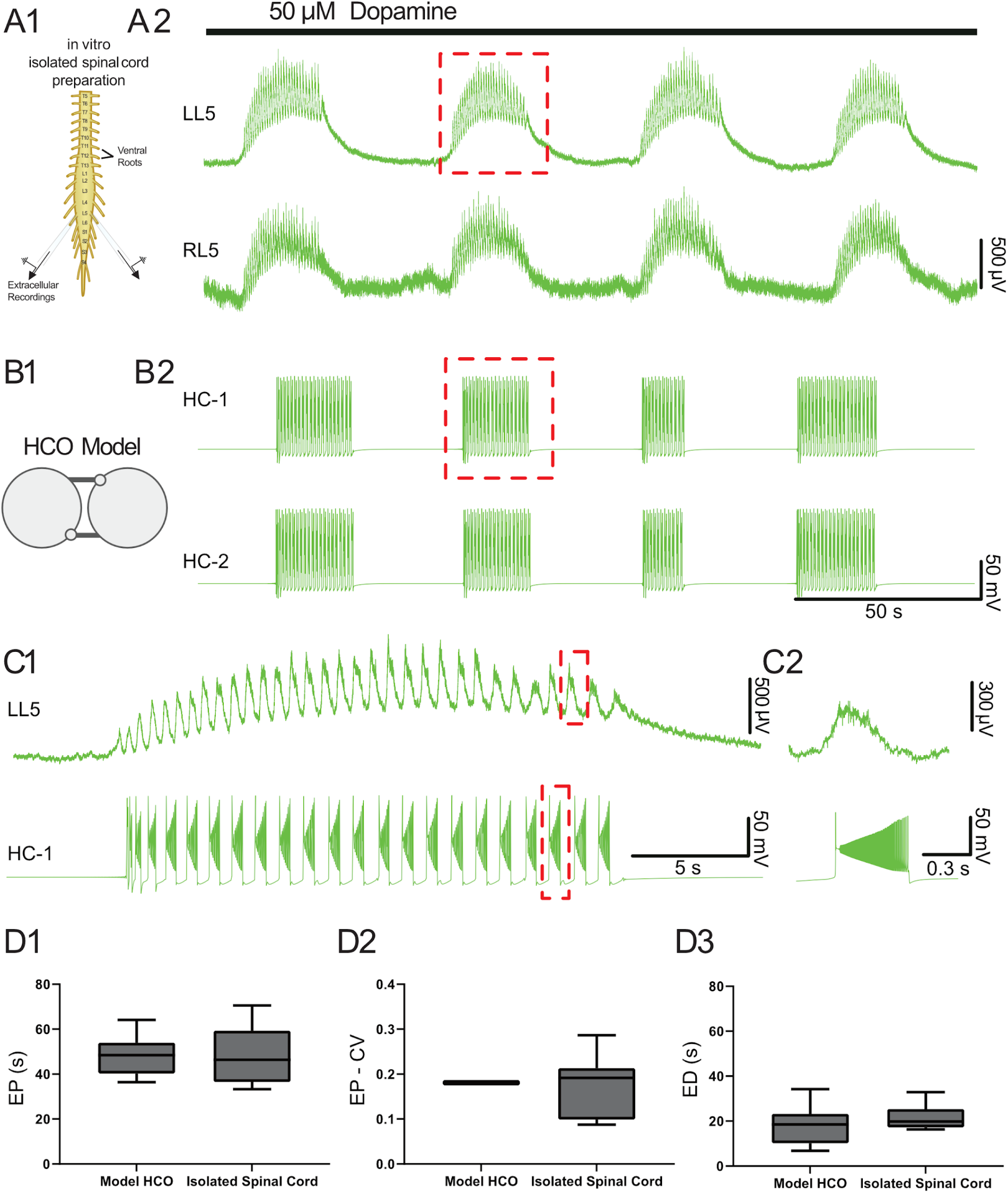
Temporal characteristics of dopamine-evoked episodic rhythmicity in the isolated spinal cord, reproduced by a biophysical model. (A1) Schematic showing recording setup used to obtain extracellular neurograms from isolated neonatal mouse spinal cords. (A2) Episodes of rhythmic activity generated by isolated spinal cords with dopamine (50 µM). Neurograms represent recordings from left and right L5 ventral roots. (B1) The half-centre oscillator (HCO) model was tuned to produce episodic rhythmicity (B2). (C1) Similar intra-episode rhythm features in a single episode of activity recorded experimentally (top trace) and from the model (lower trace) show bursting slowing down throughout the episode. (C2) The burst period and burst duration of a single intra-episode burst recorded experimentally (top trace) corresponds closely to a single burst within the model (lower trace), especially for the latter portion of the episode. (D1) Model-generated episodes (bottom trace) of activity match the recordings (top trace) in terms of episode period. (D2) episode period variability, and (D3) episode duration. Box and whisker plots depict interquartile range (boxes), median (horizontal black lines), maximum, and minimum values in data range (whiskers).

### Episodic rhythmicity depends on I_*h*_

I_h_ plays an important role in the control of rhythmic networks in both invertebrate (Peck et al. 2006; Angstadt and Calabrese 1989) and vertebrate systems (Smith and Perrier 2006; Butt, Harris-Warrick, and Kiehn 2002; Picton, Sillar, and Zhang 2018; Thoby-Brisson and Ramirez 2000; Kjaerulff and Kiehn 2001; Takahashi 1990), and dynamically interacts with I_Pump_ (Tobin and Calabrese 2005; Kueh et al. 2016; Picton, Sillar, and Zhang 2018; Zhang, Bucher, and Nadim 2017). Consistent with those findings, we found I_h_-dependent episodic bursting in both model-generated data and dopamine-elicited neurograms recorded in spinal cords in vitro. In the process of developing and validating our model, we modified parameters and equations in order to best replicate experimental data. Our initial model produced episodic and continuous rhythmicity patterns consistent with those elicited in vitro (Figure 3). However, it failed to reproduce the changes in IEI as found in our experiments with HCN-channel blocker, ZD7288 (see below). We subsequently modified the model by integrating and modulating an additional h-current (I_h-b_) while maintaining the initial h-current (I_h-a_). This resulted in a more accurate match in episodic bursting parameters and reproduced experimental results with ZD7288.

Variation of 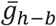 revealed two zones of parameter space that produced distinct activity patterns (episodic bursting and continuous bursting; Figure 4A, B). The model HCO switched from episodic activity to continuous bursting when 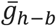 was increased beyond 0.35 nS (Figure 4B). We investigated changes in episode characteristics, including the EP, EP-CV, ED, and IEI. Within the zone supporting episodic bursting, as 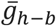 increased from 0.16 to 0.34 nS, mean ED increased from 20.2 to 180.5 s, while IEI remained roughly unchanged, mean EP increased from 50.1 s to 208 s (Figure 4B). The variability of EP represented by EP-CV significantly increased with 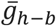 due to an increase in the variability of ED (4B1, B3). When we decreased 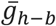 to 0 nS, episodic activity persisted with a median ED of 7.3 s, a median IEI of 32.4 s, a median EP of 40.0 s, and no change to fast bursting properties (Figure 4B, C). We compared this model parameter set with 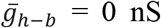 to experimental recordings where h-current was reduced by the bath application ZD7288 (Figure 4D, E).

**Figure 4:**
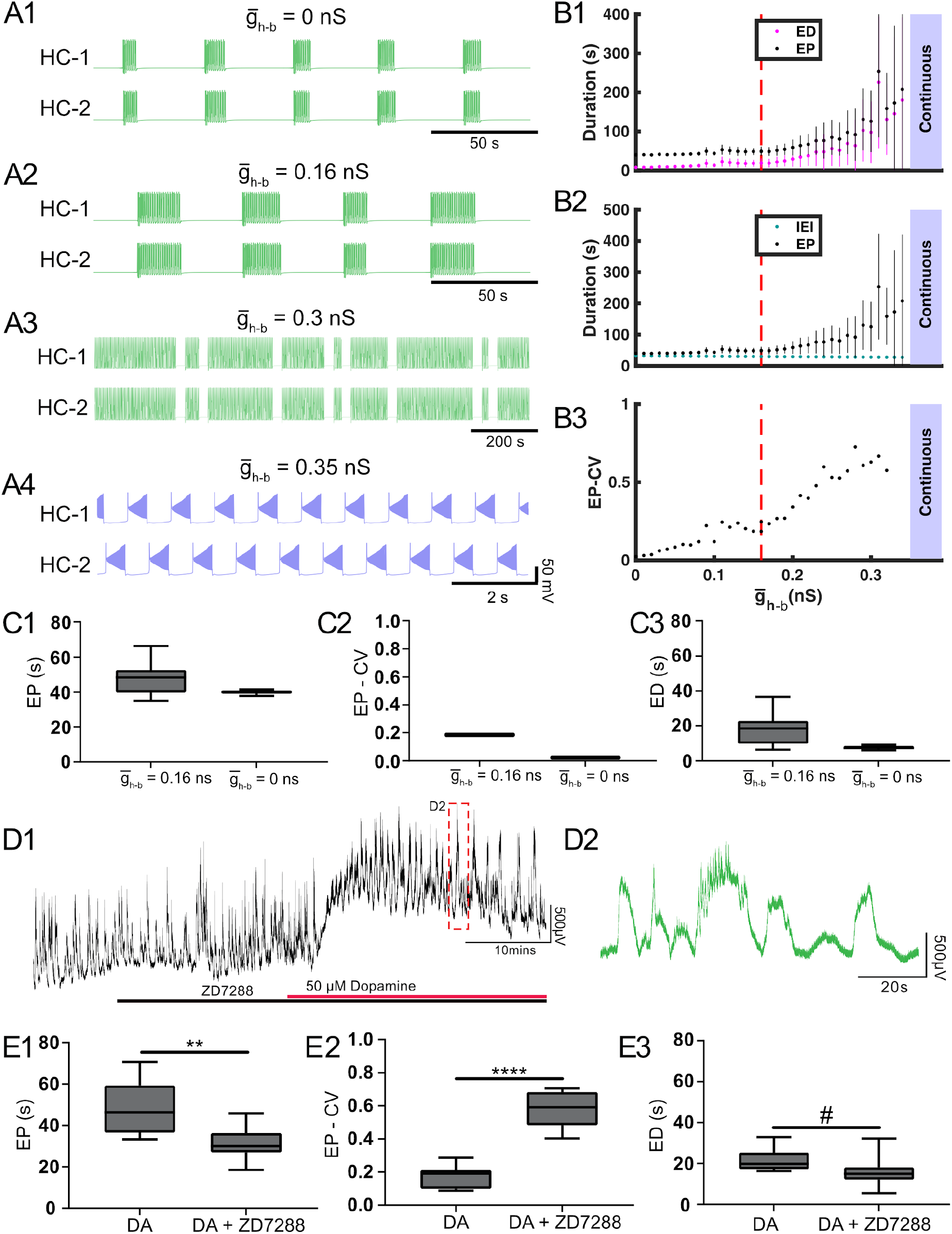
Regulation of 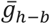 modified episodic bursting activity in the model and experimental blockade of I_h_ also affected episodic activity. (A) Examples of various types of activity produced at different values of the maximal conductance of I_h-b_ 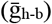 are depicted. Episodic activity occurs at 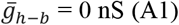, at 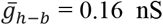 (canonical) (A2), and at 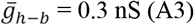, where episode duration increased with 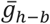. (A4) Continuous bursting occurred at 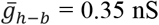. (B) Episodic activity properties and transition to continuous activity depend on 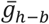. (B1) Mean episode period (EP) (black dots) and mean episode duration (ED) (magenta dots) increased exponentially as 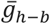 increased. (B2) Mean interepisode interval (IEI) (cyan dots) did not change significantly (p<0.05). (C) Box and whisker plots depict interquartile range (boxes), median (horizontal black lines), maximum and minimum values in the data range (whiskers). (C1) When 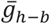 was decreased, median EP decreased, (C2) the coefficient of variation of EP (EP-CV) decreased, and (C3) median ED decreased. (D) Extracellular neurograms (D1) recorded from a single ventral root show episodic rhythmicity evoked by dopamine (black bar) in the presence of the I_h_ channel blocker ZD7288 (30–50 μM, red bar). D2 expands the inset in D1. (E) ZD7288 reduced the episode period (EP; E1), increased episode period variability (EP-CV; E2), and reduced duration (ED; E3) of episodes of dopamine-evoked rhythmicity. Box and whisker plots depict interquartile range (boxes), median (horizontal black lines), maximum, and minimum values in the data range (whiskers) for episodic rhythms evoked by dopamine alone (n = 8), or in the presence of ZD7288 (n = 11) in separate sets of experiments; asterisks indicate significance of unpaired t-tests (* p < 0.05; ** p < 0.01; *** p < 0.001; **** p < 0.0001); # indicate significance of non-parametric Mann-Whitney U test.

Consistent with our model, in isolated spinal cord preparations (n = 11), blocking I_h_ channels with ZD7288 significantly impacted episodic bursting. The direction and magnitude of change in ED and EP corresponded to the predicted effects from the model. IEIs in both experiments and modeling did not change significantly. In the presence of ZD7288, dopamine (50 µM) depolarized the ventral root neurogram (Figure 4D1), followed by the emergence of episodic rhythmic bursting (Figure 4D2). Compared to episodes elicited by preparations with dopamine alone (n = 8), dopamine-evoked episodes in the presence of ZD7288 had shorter EPs (*t*_(16)_ = 3.5, *p* = 0.003, DA: 48.5 ± 13.0; DA+ZD7288: 31.4 ± 7.9 s (mean ± SD); Figure 4E1), were more irregular (EP-CV Mann–Whitney *W* = 14, *p* = 0.02; DA: 0.17 ± 0.07; DA+ZD7288: 0.58 ± 0.1; Figure 4E2), with shorter EDs (Mann–Whitney *U* = 14, *p* = 0.02; DA: 21.5 ± 5.7, DA+ZD7288: 15.9 ± 7.0 s (mean ± SD); Figure 4E3), and no change in IEI (t_(16)_=0.5, p=0.6; DA: 17.7 ± 7.7 s; DA+ZD7288 15.7 ± 8.5 s (mean ± SD)).

We did not decrease 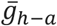 along with 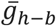 when comparing the model with the ZD7288 experimental case because the mechanism used by our model to produce episodic bursting depends heavily on the dynamics of I_h-a_ current. Due to this dependence, the model was silent when 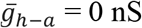 nS, while episodic activity still occurred in the ZD7288 experimental case. When 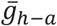 was decreased, IEI increased exponentially and then switched to silence. The extent to which hyperpolarization-activated currents are blocked by ZD7288 is dose-dependent, and the relative degree to which ZD7288 blocks each of the different types of h-current is unclear (Tanguay, Callahan, and D’Avanzo 2019; Emery, Young, and McNaughton 2012). We predict from our model that the effects of ZD7288 on episodic bursting were primarily due to a blockade of an HCN current with a lower voltage of half-activation (HCN1). Other HCN currents (HCN2 and/or HCN4) may not have been sufficiently blocked to silence the network.

### Modulation of I_Pump_ leads to transitions between episodic and continuous bursting

Four different zones of parameter space were found when I_PumpMax_ was varied (Figure 5A, B). The model exhibited episodic bursting within the range of I_PumpMax_ values between 38.5 pA and 42.5 pA (Figure 5A2-4, B). When I_PumpMax_ was decreased from that episodic regime, the model fell silent at I_PumpMax_ = 38.4 pA, and bistability of silence and continuous bursting presented when I_PumpMax_ was decreased past 37.9 pA (Figure 5A1, B). As I_PumpMax_ was decreased, the continuous bursting became more plateau-like in this bistable regime. The model fell silent again when I_PumpMax_ was decreased to 32.8 pA (Figure 5B). When I_PumpMax_ was increased from the episodic bursting regime, activity switched to another continuous bursting regime at I_PumpMax_= 42.7 pA (Figure 5A2-A5, B). Thus, the model produced episodic bursting within a range of I_PumpMax_ values and could transition from episodic bursting to continuous bursting at either end of the range (Figure 5A, B).

**Figure 5:**
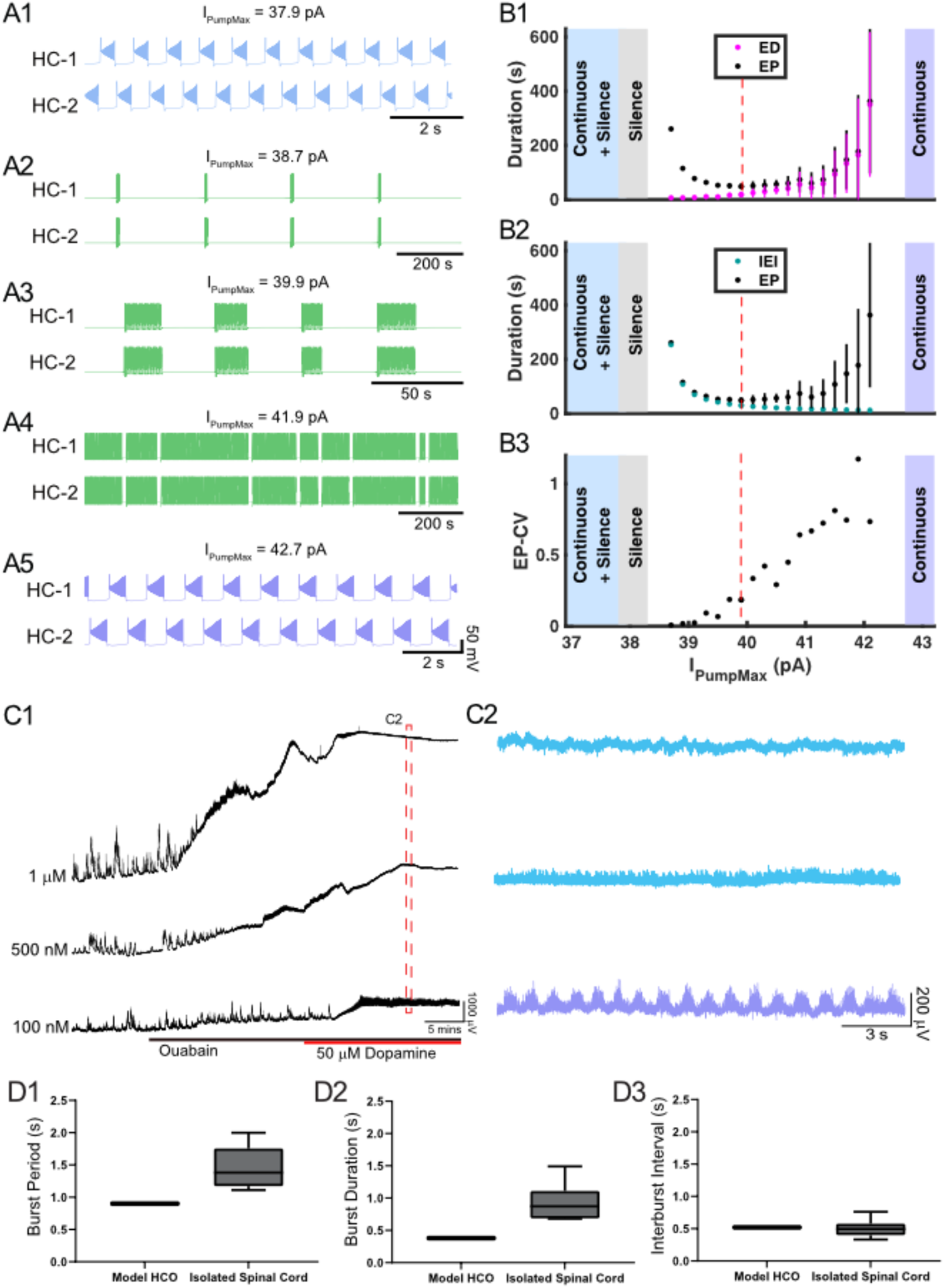
Variation in I_PumpMax_ modulates episodic bursting properties and produces transitions between episodic and continuous rhythmicity. (A) Examples of various activity types produced at different values of the maximal pump strength (I_PumpMax_) are depicted. (A1) Continuous bursting occurs at I_PumpMax_ = 37.9 pA. (A2) Episodic activity occurs at I_PumpMax_ = 38.7 pA with long IEIs and short EDs. (A3) At I_PumpMax_ = 39.9 pA (canonical), episodic activity occurs with moderately variable EDs. (A4) At I_PumpMax_ = 41.9 pA, episodic activity occurs with highly variable EDs and short IEIs. (A5) Continuous bursting occurs at I_PumpMax_ = 42.7 pA. (B) Graphs depict the relationship between I_PumpMax_ and episode parameters. (B1) As I_PumpMax_ increases from canonical, EP (black dots) and EP variability (black bars) increase due to increases in ED (magenta dots) and ED variability (magenta bars). (B2) As I_PumpMax_ decreases from canonical, EP (black dots) increases due to increased IEI (cyan dots). (B3) EP-CV increases significantly as I_PumpMax_ increases within the episodic regime. (C1) Extracellular neurograms of spontaneous activity recorded in DC from ventral roots during application of 1 µM (n = 6), 500 nM (n = 2), and 100 nM (n = 6) of ouabain (black bar) and subsequent application of dopamine (50 µM: red bar). (C2) Neurograms displaying network activity elicited by dopamine in the presence of different concentrations of ouabain highlighted in C1. (D) Box and whisker plots depict interquartile range (boxes), median (horizontal black lines), maximum and minimum values in the data range (whiskers) for burst period (BP: D1), burst duration (BD: D2), and interburst interval (IBI: D3) during continuous bursting generated by the Model HCO with I_PumpMax_ = 37.9 pA compared to continuous bursting generated by isolated spinal cords in the presence of 50 μM dopamine and 100 nM ouabain.

As I_PumpMax_ increased within the range supporting episodic bursting, ED and EP increased exponentially (Figure 5B1). The rate of change for ED and EP became particularly prominent when I_PumpMax_ was increased above 41.8 pA, where changes in I_PumpMax_ had a greater effect on ED, relative to its effect on IEI (Figure 5B1, B2). On the other hand, as I_PumpMax_ decreased toward the lower boundary between episodic activity and silent activity (38.4 pA), IEI increased exponentially, while ED decreased only slightly. The opposite trends of ED and IEI cause EP to form a U-shape (Figure 5B1, B2). The variability of EP increased significantly as I_pumpMax_ increased (Figure 5B3), mainly due to an increase in the variability of ED similar to the 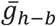 case. The continuous bursting found in our model at lower values of I_PumpMax_ is consistent with experimental results obtained when the pump current was blocked with ouabain (Figure 5D1, D2).

We validated our model’s predictions by testing the effect of varying I_Pump_ strength using different concentrations of ouabain on the rhythmic activity in the isolated spinal cord. Ouabain has dose-dependent effects on I_Pump_, preferentially disrupting the ɑ_3_ subunit and dynamic pump function at low concentrations while impairing the α_1_ subunit at higher concentrations (Lichtstein and Rosen 2001). Prior to application of dopamine, ouabain concentrations of 1 µM (Figure 5C; n = 6; mean ± SD Δ DC potential = 3433 ± 1897 µV) and 500 nM (n = 2; mean ± SD Δ DC potential = 738 ± 128 µV) dose-dependently depolarized ventral root DC potentials (Figure 5C1, B1: *F*_(2,11)_ = 9.7, *p* = 0.004) and suppressed spontaneous network activity (Figure 5C1: *F*_(2,11)_ = 10.5, p = 0.003). Subsequent application of dopamine when 500 nM or 1 µM of ouabain was present further depolarized the ventral roots but did not elicit any superimposed rhythmicity (Figure 5C2; top two traces in blue), consistent with the silent state predicted by the model at the lowest I_PumpMax_ strengths. As predicted by the model, reducing ouabain concentration to 100 nM produced continuous rhythmic bursting in the presence of dopamine with burst metrics within a range as those generated by the model (Figure 5C1, C2). Continuous bursting elicited by dopamine in the presence of 100 nM of ouabain had a mean cycle period (CP) of 1.45 ± 0.37 s, a CP-CV of 0.22 ± 0.07, burst duration of 0.93 ± 0.33 s, IBI of 0.50 ± 0.2 s, and were within range of the burst characteristics generated by the HCO model with I_PumpMax_ = 37.9 pA (Figure 5 D1-D3). Episodic bursting returned in three of six preparations following a wash with 50 µM dopamine in regular aCSF (not shown).

We next set out to upregulate I_Pump_ with monensin, an antibiotic that acts as a Na^+^-H^+^ antiporter, leading to increased [Na]_i,_ indirectly increasing I_Pump_ activity (Kueh et al. 2016; Picton et al. 2017). Prior to application of dopamine, 2 µM monensin (n = 12) reduced spontaneous bursting activity. Subsequent application of 50 µM dopamine with monensin depolarized the ventral root potentials and produced episodic bursting in 10 preparations (Figure 6A). In 2 preparations we observed a depolarization following DA application, but no episodic rhythmic bursting was evoked). Episodes evoked by dopamine in the presence of monensin (n=10) had a longer EP (t_(16)_ = 2.3, p = 0.03), higher EP-CV (t_(16)_ = 9.2, p = 8.2e-8), and longer IEI (t_(16)_ = 4.5, p = 0.0004), with no change in ED (t_(16)_ = 0.8, p = 0.4) (Figure 6B). Consistent with experimental findings, applying ‘monensin’ in the HCO model, represented by a monensin rate constant M = 0.0025, led to an increase in EP and IEI, no change in ED, but decreased EP-CV (Figure 6C).

**Figure 6:**
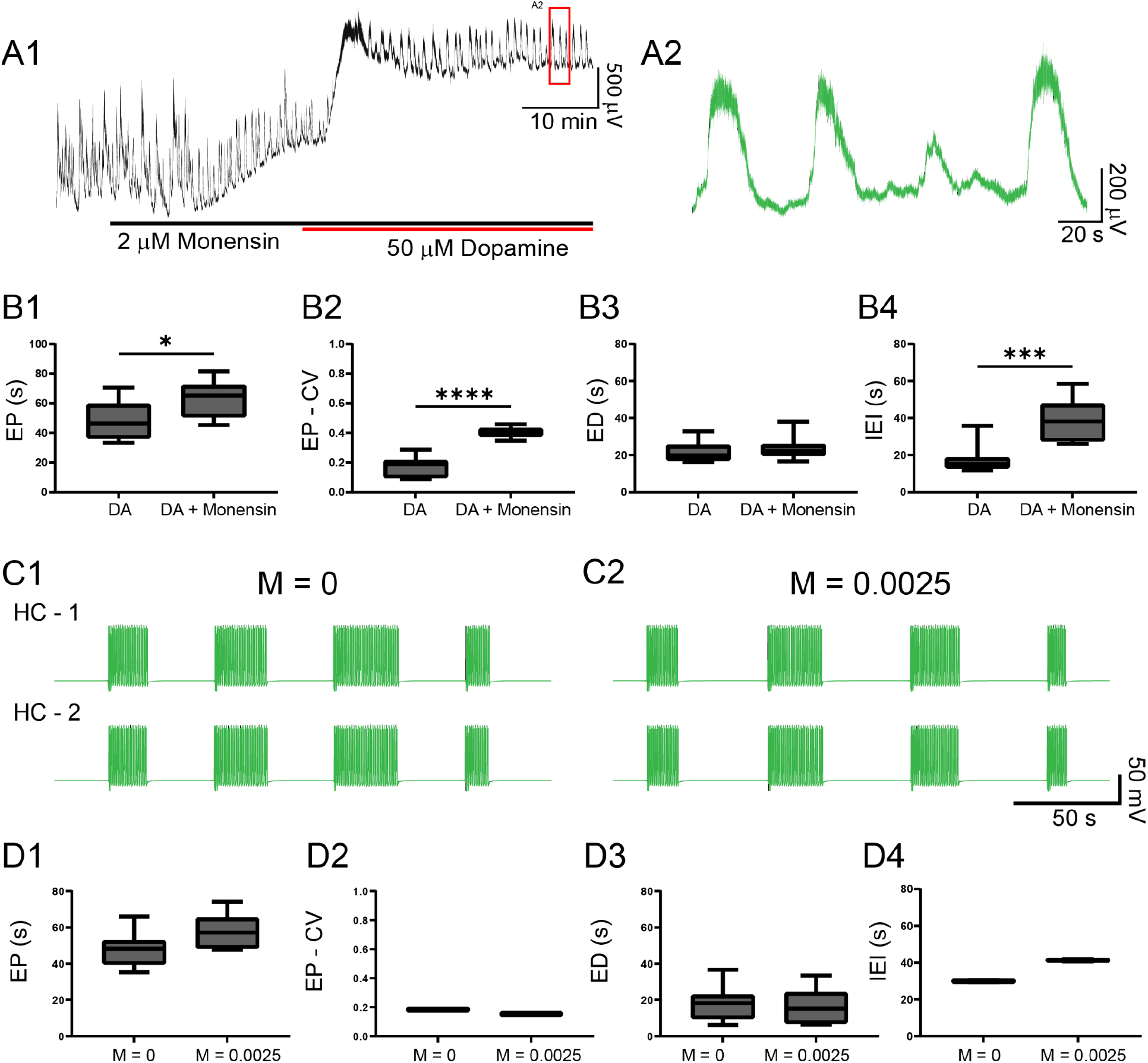
Monensin increases episode period and IEI in both modeled and experimental data. (A1 Extracellular neurograms recorded from ventral roots with bath application of 2 µM monensin (black bar along the x-axis) followed by 50 µM dopamine (red bar along x-axis). A2)Expanded segment from inset (red box) in A1. (B1) Episode period (EP) increases with monensin (B2) but becomes more variable, (B3) while ED is maintained, and (B4) interepisode interval is increased. Asterisks denote significance level following unpaired t-tests (DA: n = 8; DA+ Monensin: n = 10). We simulated monensin application in the model HCO by increasing the influx of intracellular sodium [Na]_i_ using a monensin rate constant (M). Changing this rate constant from M = 0 (C1) to M = 0.0025 (C2) in the model reproduced most of the effects of monensin on isolated spinal cords, leading to (D1) an increase in EP, (D2) decrease in EP-CV, (D3) no change in ED, and (D4) an increase in IEI.

## Discussion

Spinal circuits produce a diverse array of rhythmic motor outputs, a subset that can be generated at birth (Whelan, Bonnot, and O’Donovan 2000). Episodic activity has been reported in developing spinal circuits for a range of species (Picton, Sillar, and Zhang 2018; Combes et al. 2004; Mahrous and Elbasiouny 2018; Kondratskaya et al. 2019; Wiggin et al. 2012; McDearmid and Drapeau 2006; Gozal et al. 2014; Sharples and Whelan 2017). Episodic and continuous patterns of rhythmic activity can be generated using the in vitro perinatal mouse spinal cord, and transitions between patterns can be induced by manipulating neuromodulatory tone or excitability (Sharples and Whelan 2017). Here, we focused on cellular mechanisms that produce episodic and continuous patterns of rhythmic activity and the mechanisms that govern transitions between them (Figure 7). We developed a biophysical model that produces episodic and continuous patterns of bursting activity and investigated how key network and cellular properties contribute to their generation in isolated spinal cords. The model was a single CPG composed of two inhibitory half centres that could adjust parameters of episodic bursting and switch output modes through modulation of I_Pump_ and I_h_. We derived these parameters from insights based on studies on the dynamic roles of I_Pump_ and I_h_ in the regulation of the leech heartbeat (Tobin and Calabrese 2005; Kueh et al. 2016) and *Xenopus* tadpole locomotor CPGs (Zhang and Sillar 2012; Zhang et al. 2015; Picton, Sillar, and Zhang 2018).

**Figure 7:**
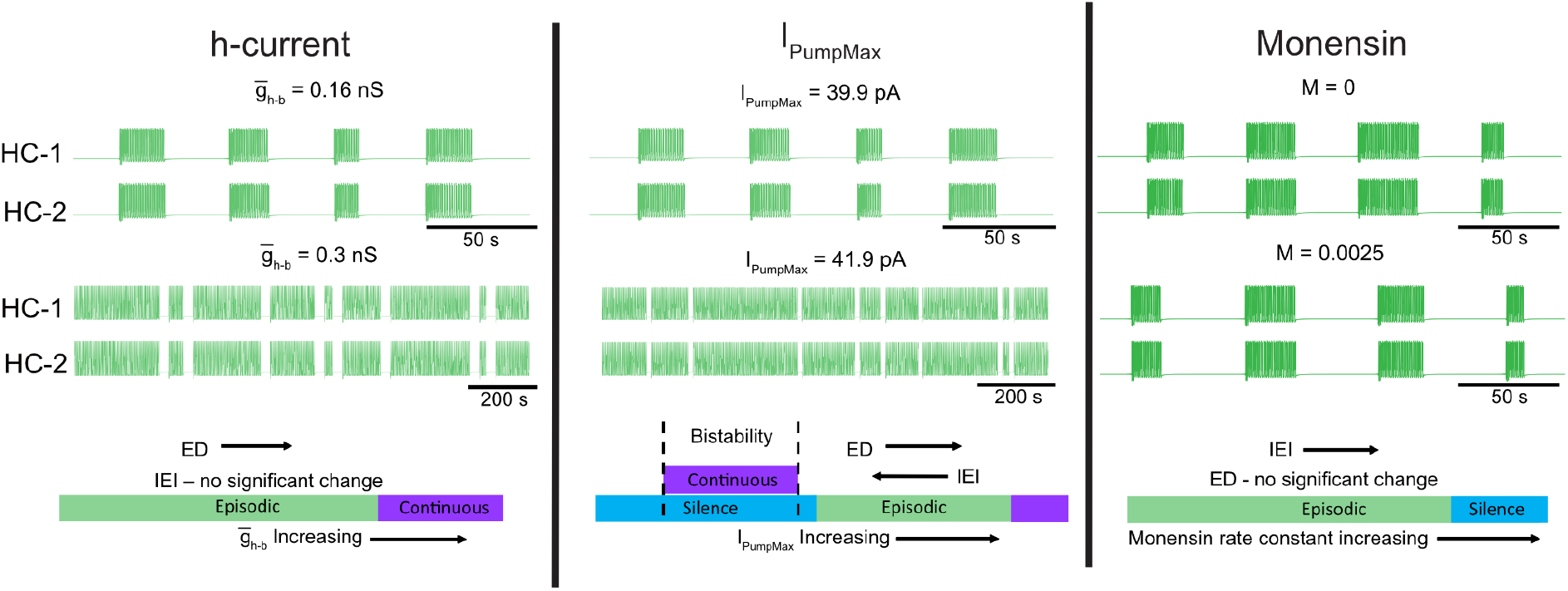
Neuromodulators evoke diverse rhythmic outputs by moving the network through an excitation-based parameter space by altering I_PumpMax_, and 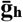. Left panel represents two traces showing behavior of the model while altering the h-current. Middle panel illustrates the changes in episodic activity following manipulation of I_PumpMax_ and the right panel illustrates model output before and after the introduction of a monensin factor. The bottom schematics summarize the changes in rhythmic characteristics following manipulation of the h-current and I_PumpMax_, or through introduction of a monensin factor.

Of the two h-currents in our model, one was necessary for episodic bursting, and the other modulated episode duration leading to a transition to continuous bursting when increased. Consistent with this finding, blocking I_h_ reduced episodic bursting duration evoked by dopamine in vitro (Figure 3). In the case of I_Pump_, the model revealed that either up or down-regulation of I_PumpMax_ produced a transition to continuous bursting, a finding supported by our in vitro pharmacological experiments (Figure 4). Our results demonstrate that multiple pathways can lead to transitions between episodic and continuous rhythmic patterns (Figure 7). This finding is consistent with studies modelling the stomatogastric ganglion (STG), which demonstrated that disparate circuit properties produced similar outputs (Prinz, Bucher, and Marder 2004) and led to transitions between circuit outputs (Gutierrez and Marder 2014; Gutierrez, O’Leary, and Marder 2013).

### Dynamic mechanisms lead to transitions between episodic and continuous rhythmic activity in a single developing CPG

We provide an analysis of cellular dynamics when each element in our model is individually manipulated. We find parallel changes in biological network behaviour when these elements are individually modulated pharmacologically. The pump-mediated ultraslow afterhyperpolarization (usAHP) is expressed in mouse spinal interneurons and motoneurons at early postnatal stages, and is modulated by dopamine (Picton et al. 2017). Of the properties established in our model, we found that I_Pump_ plays a critical role in generating episodic bursting with manipulations in I_PumpMax_ leading to switching between episodic and continuous bursting through modulation of the IEI. In line with our model, blocking the pump with low concentrations of ouabain (100 nM), which may select toward blocking the α_3_ subunit of the pump (Lichtstein and Rosen, 2001), promotes the transition from episodic to continuous bursting. In contrast, higher concentrations of ouabain (0.5–1 µM) inhibited rhythmic activity in the presence of dopamine and is consistent with the silent output predicted by the model at the lowest I_PumpMax_ strengths. Manipulating the pump by increasing intracellular sodium experimentally with monensin caused an increase in EP and IEI while ED remained constant. This is consistent with modelled data where I_Pump_ was increased indirectly through the introduction of a ‘monensin factor’.

Interestingly, increasing I_PumpMax_ in the model also prompted a transition from episodic to continuous bursting. Increased pump activity may produce transitions from episodic to continuous locomotor behaviours observed during development in fish and tadpoles (Lambert, Bonkowsky, and Masino 2012; Buss and Drapeau 2001; Müller, Stamhuis, and Videler 2000; Gabriel et al. 2011; Hachoumi and Sillar 2020). However, our work raises the possibility that a reduction in dynamic pump function may also generate such transitions. Thus, modulating the pump’s function may change the mode of operation during development (Gonzalez-Islas, Garcia-Bereguiain, and Wenner 2020), in addition to dynamically regulating locomotor behaviour.

Rhythm-generating interneurons also often express the mixed cation-conducting current I_h_ (Borowska et al. 2013; Dougherty and Kiehn 2010; Wilson et al. 2005; Kiehn et al. 2000), which is activated at hyperpolarized potentials and supports escape from inhibition, which in turn facilitates burst initiation. Similarly, in the *Xenopus* tadpole spinal cord CPG, I_h_ is expressed only in glutamatergic rhythm-generating interneurons (dINs), where it opposes and mitigates the inhibitory influence of I_Pump_ on the membrane potential (Picton, Sillar, and Zhang 2018). I_Pump_ also interacts with I_h_ in leech heartbeat CPG neurons, which accounts for the paradoxical speeding up of the rhythm when I_Pump_ is stimulated experimentally with monensin (Kueh et al. 2016). Our data is consistent with this; we found that increasing both I_PumpMax_ and I_h_ led to a transition from episodic to continuous rhythms. Notably, I_h_ plays an important role in controlling the ED, which is supported by both our model and experimental results, and also episode stability which is consistent with our previous work demonstrating a prominent role for dopamine in reducing variability in the cycle period in locomotor-like rhythms (Sharples et al. 2015). The bidirectional receptor-dependent modulatory effects of dopamine (Sharples et al. 2020) could be mediated by adjusting the balance of the I_Pump_ (Picton et al. 2017) and I_h_ in spinal neurons.

Given that there was no change in the IEI elicited by ZD7288, a reduction in EP may be due to network compensation to maintain phase relationships in response to a reduction in ED. Indeed, EP variability is increased by ZD7288, producing network output that is consistent with spontaneous activity generated in the absence of stimulation (Dalrymple et al. 2019), supporting the notion that I_h_ is essential for the generation of robust episodic activity. It is unclear whether ZD7288 selectively or uniformly targets each of the HCN subunit isoform channels (Emery, Young, and McNaughton 2012; Tanguay, Callahan, and D’Avanzo 2019); therefore it is possible that different HCN channels and their selective modulation may subserve dedicated functions in the control of spinal circuits (Kjaerulff and Kiehn 2001). Alternatively, ZD7288 has been reported to block other channels including Cav1.3 and NaV1.4 (Sánchez-Alonso, Halliwell, and Colino 2008; Wu et al. 2012) which contribute to T-type calcium currents and persistent sodium currents respectively, and could also contribute to the effects on network stability that were not predicted by our model.

Our data support multiple mechanisms that lead to transitions between episodic and continuous bursting patterns with modulation of I_h_ adjusting the ED and I_Pump_ controlling the IEI. This work extends the knowledge on the pump’s diverse functions in the dynamic regulation of neuronal excitability described in a wide range of model systems (Pulver and Griffith, 2010; Zhang, Bucher, and Nadim 2017; Zhang and Sillar 2012; Picton, Sillar, and Zhang 2018; Ballerini, Bracci, and Nistri 1997; Gerkau et al. 2019; Tiwari et al. 2018; Kueh et al. 2016; Picton et al. 2017). Given that I_Pump_ and I_h_ are important for the dynamic activity-dependent regulation of neural networks, and in some cases, producing short term motor memory (STMM) (Picton et al., 2018), an interesting area for future investigation would be to determine how modulation of these properties can adjust network behaviours such as STMM (as discussed by (Hachoumi and Sillar 2020)).

### Flexible outputs produced by a single multifunctional CPG

An important question that remains only partially resolved is whether spinal circuits produce diverse rhythmic activities by recruiting multiple dedicated rhythm generating circuits or through the modulation of a single multifunctional CPG? The latter appears to be true in many vertebrate systems and has been well described in invertebrate CPGs (Blitz et al. 1999; Briggman and Kristan 2006). For example, in the turtle, scratching and swimming movements are produced by a combination of overlapping and dedicated populations of spinal interneurons (Hao et al. 2011; Berkowitz 2002; Snyder and Rubin 2015). Similar results have been found for swimming and struggling motor patterns in both *Xenopus* tadpoles (Soffe 1993; Li et al. 2007) and larval zebrafish (Liao and Fetcho 2008). Overlapping populations of spinal interneurons appear to contribute to the generation of walking and paw-shaking in cats (Barbeau and Rossignol 1990; Carter and Smith 1986), and our previous computational models proposed a multifunctional CPG with dedicated roles for calcium and sodium conductances in the generation of these two rhythmic motor patterns (Parker et al. 2018; Parker, Khwaja, and Cymbalyuk 2019). The episodic and continuous bursting modes that we describe here could serve as a new system to study how mammalian spinal circuits produce diverse rhythmic activities. Whether these two patterns are produced by dedicated or overlapping neural elements in the spinal cord remains to be determined.

Dialogue between experimental and computational systems offers insight into mechanisms of multifunctionality. Computational models inspired by the organization of the STG have elegantly demonstrated that degenerate mechanisms in electrical and chemical inhibitory synapses can produce pattern switching in a small multifunctional circuit (Gutierrez, O’Leary, and Marder 2013). Our computational model, composed of an inhibitory HCO, suggests that modulation of key intrinsic properties can lead to multifunctionality within a single circuit. Indeed our HCO model represents a fundamental element of current computational models of spinal CPGs (Ausborn et al. 2018; Yakovenko et al. 2005; Shevtsova et al. 2020; Rybak, Stecina, et al. 2006) and may serve as a key locus to produce flexible rhythmic outputs. Therefore, this work offers insight into how spinal circuits and computational models of locomotor CPGs can produce adaptable locomotor output with the integration of key intrinsic properties as targets for neuromodulators serving as a basis for their control. Our focus here is on the generation of episodic locomotor activity which contains both slow episodic and fast locomotor components and we show that a simplified HCO can produce both elements. While we show that I_h_ and I_Pump_ in an inhibitory circuit are key properties for episodic and continuous rhythms, there are likely others.

### Limitations of the model

Our model is a basic HCO consisting of two units with mutual inhibitory synaptic connectivity. We realize that the vertebrate spinal locomotor circuit is more complex with recurrently connected glutamatergic interneurons. In addition, interposed inhibitory interneurons that are part of the spinal locomotor circuit are not included in our model. Rhythmic activities generated by the spinal cord or computational models are often glutamate-dependent (Buchanan and Grillner 1987; Dale and Roberts 1984; Talpalar and Kiehn 2010; Masino et al. 2012) with persistent sodium currents (Song et al. 2020; Del Negro et al. 2002; Brocard et al. 2013; Tazerart, Vinay, and Brocard 2008; Verneuil et al. 2020; Shevtsova et al. 2020) and electrical coupling (Shevtsova et al. 2020; Dougherty et al. 2013; Ha and Dougherty 2018; Wilson et al. 2005; Hinckley and Ziskind-Conhaim 2006; Hinckley et al. 2005) being particularly important for sustaining rhythmic activity. While our model included a persistent sodium current to support intrinsic bursting, the cell types, in addition to synaptic and intrinsic mechanisms that *generate* episodic bursting, remain to be systematically investigated. That said, similar theoretical modelling approaches have been implemented, and reduced HCO models of locomotion similar to ours mimic the behavior of more complex population-based models (Ausborn et al. 2018; Shevtsova et al., 2020).

### Future directions

Episodic swimming in larval zebrafish is generated by a distributed population of glutamatergic spinal neurons (Wiggin et al. 2012; Wiggin, Peck, and Masino 2014; Wahlstrom-Helgren et al. 2019) and episodic rhythms elicited in isolated rat spinal cords are diminished by blockers of glutamatergic transmission (Marchetti and Nistri 2001). In support of those findings, our previous work demonstrated a role for NMDA receptors in maintaining episodic rhythms and enabling transitions to continuous rhythms (Sharples and Whelan 2017). Future work adapting our model to include mutually excitatory synapses will shed light on the underlying mechanisms that contribute to episode initiation. Furthermore, complementary experimental approaches harnessing genetic tools available to study mice and zebrafish will also offer insight into putative “episode-generating” spinal interneurons. V3 interneurons are one population of interest, given their intrinsic and synaptic properties, and their role in supporting locomotor bouts (Zhang et al. 2008; Chopek et al. 2018; Danner et al. 2019). Alternatively, episodic bursting may be supported by astrocytes that have broad, distributed connectivity, regulate spinal motor circuits (Broadhead and Miles 2020; Carlsen and Perrier 2014; Acton, Broadhead, and Miles 2018; Carlsen et al. 2021), have well-established roles in regulating glutamatergic transmission at tripartite synapses (Perea, Navarrete, and Araque 2009), and form complex synaptic structures within the newborn and adult mouse spinal cord (Broadhead et al. 2020).

## Conclusions

Our modelling and in vitro experiments reveal that a single CPG could produce episodic activity and intra-episode bursting, and that degenerate mechanisms lead to transitions between silence, episodic, and continuous bursting through modulation of I_Pump_ or I_h_. While episodic and continuous patterns have been well documented in isolated developing mammalian spinal circuits (Barbieri and Nistri 2001; Marchetti and Nistri 2001; Gozal et al. 2014; Sharples and Whelan 2017), these activities may not be restricted to developing nervous systems; there are examples of these different patterns occurring in adult fish, rodents, and cats (Crawley 2007; Wiltschko et al. 2015; Gabriel et al. 2011; Müller, Stamhuis, and Videler 2000; Mahrous and Elbasiouny 2018). This study provides insight into the network mechanisms that govern the generation of context-dependent locomotor patterns produced during different states in freely behaving animals and in developing motor networks.

## Acknowledgements

We acknowledge support from the Whelan Lab. We also thank Dr.’s Tuan Bui, Ronald Calabrese, and Keith Sillar for helpful comments on an early version of the manuscript.

## Contributions

SAS, APL, NC, and AS performed in vitro experiments and SAS analyzed the data with code written by LY. The computational model was developed and analyzed by AV, JP, and GSC. SAS, AV, JP., GSC, and PJW prepared figures and wrote the manuscript. All authors approved the final version of the manuscript. SAS, GSC, and PJW conceived and designed the research.

## Methods

### Ethical approval & animals

Experiments were performed on neonatal C57BL/6 (n = 45) mice, postnatal 0–4 days old (P0– P4). The University of Calgary Health Sciences Animal Care Committee approved all procedures (protocol number AC16-0182).

### Tissue preparation

We anesthetized the animals via hypothermia by placing them at −20 °C for 5–10 minutes. Animals were then decapitated, eviscerated, and the spinal column was pinned ventral side up in a dissecting dish lined with silicone elastomer (Sylgard, MilliporeSigma, Oakville, ON, Canada) and superfused with room temperature (21 ^o^C) carbogenated (95% O_2_-5% CO_2_) artificial cerebrospinal fluid (aCSF; in mM, 128 NaCl, 4 KCl, 1 MgSO_4_, 1.5 CaCl_2_, 0.5 Na_2_HPO_4_, 21 NaHCO_3_, 30 D-glucose; total osmolarity: 310–315 mOsm). The spinal cord was isolated by performing a ventral laminectomy. The nerve roots connecting the spinal cord to the vertebral column were subsequently cut, isolating the spinal cord. We then transferred the isolated spinal cord to a recording chamber perfused with carbogenated aCSF and placed it ventral side up. We gradually increased the bath temperature to 27 °C (Whelan et al., 2000). The advantage of this method is that it is close to physiological temperature and avoids fluctuations in room temperature. We left the spinal cords to stabilize for 1 hour before performing experiments.

### Electrophysiology

Extracellular electrophysiological recordings were obtained from ventral roots with tight-fitting suction electrodes fashioned from polyethylene tubing (PE50). Total signal amplification was 1000×, including a pre-amplifier (10×) and second stage amplifiers (Cornerstone EX4-400 Quad Differential Amplifier) at 100×. Amplified signals were acquired in DC and digitized at 2.5 KHz (Digidata 1440A/1550B; Molecular Devices; Sunnyvale, CA). We saved data acquired using Clampex 10.7 software (Molecular Devices) to a desktop computer for offline analysis.

### Pharmacology

Episodic rhythmicity was evoked by bath application of dopamine hydrochloride (50 µM; Sigma-Aldrich). We blocked the I_Pump_ with ouabain (100 nM-1 µM; Tocris), and I_h_ - producing hyperpolarization-activated cyclic nucleotide (HCN) channels with ZD 7288 (30–50 µM; Tocris). I_Pump_ was potentiated by monensin (monensin sodium salt, 2 µM dissolved in ethanol (0.03%), Sigma, M5273). All pharmacological agents were prepared following solubility guidelines specified by their respective vendors. We were careful to ensure that volumes of drugs prepared in dimethyl sulfoxide (DMSO) did not exceed 0.04% (vol/vol) concentration in working solutions.

### Computational model

The model consists of two identical and mutually inhibitory neurons, which represents one side, flexor and extensor, of a spinal CPG, assembled as an HCO. Each neuron, described by a single equipotential compartment, represents the activity of a population (flexor or extensor) of controlling interneurons. Each neuron is equipped with ionic currents modelled using Hodgkin-Huxley formalism: slowly inactivating persistent Na^+^ current (I_NaP_), low-threshold slowly inactivating Ca^2+^ current (I_CaS_), a fast Na^+^ current (I_NaF_), a delayed rectifier-like K^+^ current (I_KDR_), a fast transient K^+^ current (I_KA_), an inhibitory spike-mediated synaptic current (I_Syn_), and two types of hyperpolarization-activated inward (h-) current, I_h-a_ and I_h-b_. The two h-currents differ mainly by their voltages of half-activation, where V_1/2h-b_ is much more hyperpolarized than V_1/2h-a_. The fast K^+^ current, I_KA_, was implemented using experimental data from previous models (Hayes et al. 2008). It’s activation variable is fast and is implemented as instantaneous. The equations for I_NaF_ and I_KDR_ were obtained from earlier work (Rybak, Shevtsova, et al. 2006) and implemented along with I_CaS_, I_Syn_, and I_NaP_ described in (Bondy et al. 2016). Our descriptions of the h-currents and leak currents were adapted from our previous study (Kueh et al. 2016) and had corresponding reversal potentials for Na^+^ and K^+^ components. The two components of the h-currents shared the same conductance and activation values, but conductance was scaled differently, with 3/7 and 4/7 for the Na^+^ and K^+^ components, respectively. The ratio of the Na^+^ component to the K^+^ component was determined using a reference reversal potential, E_LRef_.

The membrane potential of each neuron in the HCO is governed by the following current conservation equation:

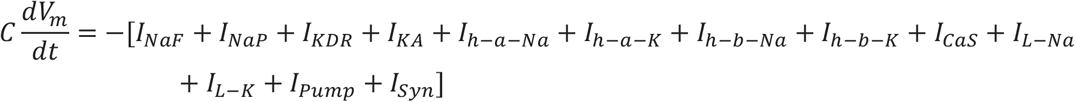

where C is the membrane capacitance in nF, *V*_*m*_ is the membrane potential in mV, and t is time in seconds, and *I*_*xyz*_ are ionic membrane currents: They are described by the following formula:

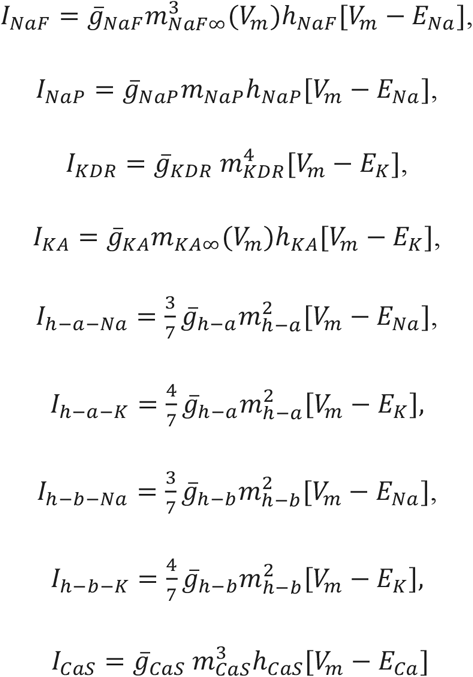

and two components of the leak current:

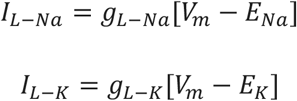

where leak conductances were computed with the reference values of equivalent total leak conductance *g*_*L*_ and reversal potential *E*_*LRef*_ using the Na^+^ reversal potential:

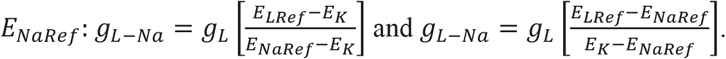

The description of the pump current was adapted from (Cressman et al. 2009); it has maximal value *I*_*PumpMax*_ and is activated by [Na]_i_ and [K]_o_ with concentrations of half-activation 25 mM and 6 mM, respectively:

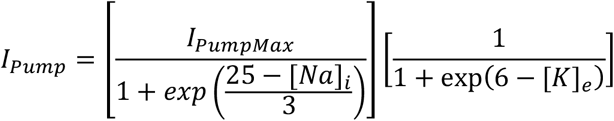

*I*_*Syn*_ is the synaptic current:

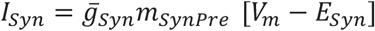

where *m*_*SynPre*_ is a synaptic activation variable that depends on the membrane potential of the presynaptic cell. The gating variables are mostly governed by equations using the following Boltzmann function: 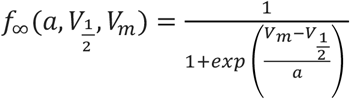

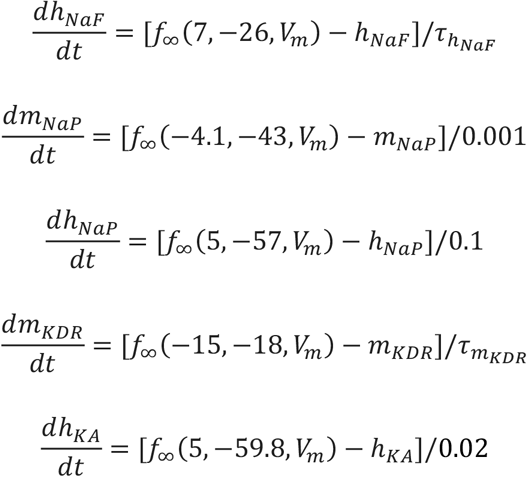

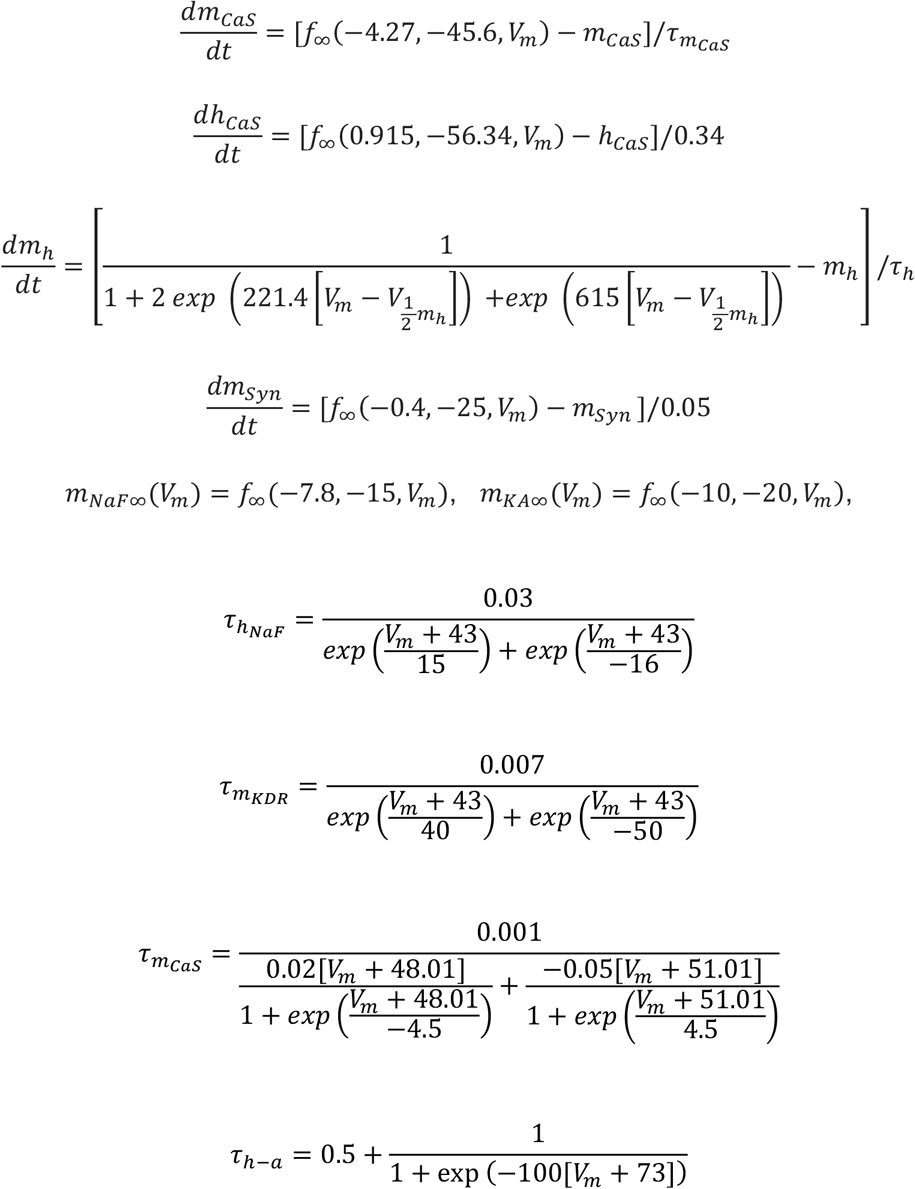

Intracellular potassium concentrations ([K]_i_) and extracellular potassium concentration ([K]_o_) are fixed parameters. Our model has dynamic intracellular Na^+^ concentration ([Na]_i_) and fixed extracellular Na^+^ concentration ([Na]_o_). To compute [Na]_i_, the model accounts for Na^+^ influx carried by the Na^+^ currents (I_NaP_ and I_NaF_) and Na^+^ components of leak (*I*_*L*−*Na*_) and h- (*I*_*h*−*Na*_) currents, and outflux produced by the Na^+^/K^+^ ATPase pump current (I_Pump_):

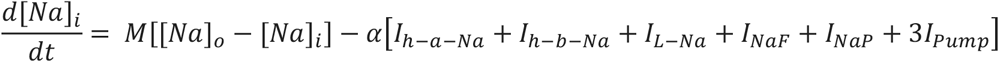

where 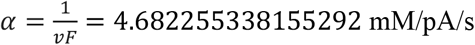, v is intracellular volume and F is the Faraday’s constant. We simulate the application of monensin by increasing the monensin rate constant (M) from zero. This rate constant is M = 0 unless stated otherwise.

Reversal potentials for Na^+^ and K^+^ currents are computed as the Nernst potentials 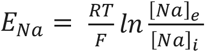 and 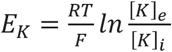 with 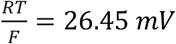, using fixed [*Na*]_*e*_ = 120 *mM*, computed state variable [*Na*]_*i*_, and fixed [*k*]_*e*_ = 9 *mM* and [*k*]_*i*_ = 130 *mM*. Canonical values of other model parameters are *C* = *0*.*001 nF*, 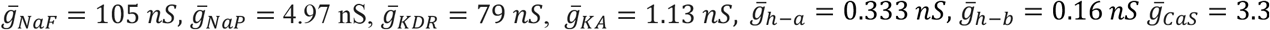 *nS, E*_*Ca*_ = 160 *mV, g*_*L*_ = 1.8816 *nS, E*_*LRef*_ = −55 *mV, E*_*NaRef*_ = 65 *mV*, 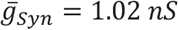, *E*_*syn*_ = −70 *mV*.

### Data analysis

The model was implemented in C programming language and analyzed data using MATLAB (Mathworks Inc, Natick, MA.). Integration was performed using the explicit embedded Runge-Kutta Prince-Dormand (8,9) method with an absolute error tolerance of 10^−8^, relative error tolerance of 10^−9^, and an initial step size of 10^−8^ (the GNU scientific library; (Galassi et al. 2002). The model was integrated at each parameter value investigated for 5000 seconds before analyzing to ensure that the model had reached its stable state.

Episodes of rhythmicity were analyzed using custom-written MATLAB scripts to detect episode onset, offset, and amplitude. We determined episode onset and offset from DC signals that were detrended, band-pass filtered (0.01–1 Hz), and smoothed with a Gaussian-weighted moving average. Peak amplitude of each detected episode was extracted from raw, unprocessed signal. We then calculated EP, ED, lag time, and phase in Excel from episode onset and offset times. The EP coefficient of variation (EP-CV) was set to distinguish regular (> 0.2) and irregular (< 0.2) patterns of episodic bursting.

### Statistical analysis

We analyzed changes in characteristics of episodic rhythmicity recorded in ventral roots over time using one-way or two-way repeated measures analyses of variance (ANOVA), as appropriate. Pharmacological manipulations of spinal circuits were compared to appropriate time-matched preparations using unpaired t-tests. Data that violated assumptions of normality (Shapiro–Wilk test) or equal variance (Brown–Forsythe test) were analyzed via nonparametric Mann–Whitney *U* (if two groups) or Kruskal– Wallis (if more than two groups) tests. All effects surpassing a significance threshold of *p* < 0.05 were further examined with Holm-Sidak post hoc tests to compare all treatment conditions to the appropriate normalized time-matched vehicle control.

